# Experimental evolution of promiscuous kin recognition from a homotypic specific cell surface receptor

**DOI:** 10.1101/2025.05.04.652116

**Authors:** Tingting Guo, Daniel Wall

## Abstract

Recognizing the difference between self and nonself is a crucial step in the development of multicellularity. *Myxococcus xanthus* is a model organism for studying these processes during the transition from single cell to multicellular life. The polymorphic cell surface receptor TraA helps to mediate these transitions by directing cooperative behavior toward kin or clonemates. TraA is a highly specific receptor, capable of recognizing other TraA proteins with identical or nearly identical sequences by homotypic binding, but the molecular basis of recognition is poorly understood. In this study, we designed a targeted TraA library, consisting of thousands of variants, which changed 10 predicted specificity residues. By screening this library, we identified TraA variants with different combinations of substitutions that resulted in altered recognition, often leading to promiscuous TraA-TraA binding. Additionally, we identified key residues that dictate specificity between distant TraA groups and showed that changing these residues altered the recognition specificity. Furthermore, we propose a model to explain how TraA recognition specificity evolved through the generation of intermediate promiscuous variants driven by reward/punishment interactions. Our results highlight the malleable nature of the TraA variable domain involved in specificity, shedding light on the molecular and evolutionary basis of social recognition in *M. xanthus*.

## Introduction

The motile soil bacterium *Myxococcus xanthus* is renowned for its complex social behaviors. Its lifestyle involves transitions from solitary cells to multicellular cooperative tissues, which include self-recognition during aggregation, where cooperative behaviors are directed toward clonemates or close kin. To achieve this, cells rely on kin recognition and discrimination systems. One such mechanism is outer membrane exchange (OME), which enables *M. xanthus* to distinguish between self and nonself. During OME, cells exchange substantial amounts of outer membrane (OM) proteins, lipids, and lipopolysaccharides through direct cell-cell contact, thought to be mediated by transient OM fusion [1–3].

OME governs diametrically opposed functions, fostering both cooperation and antagonism among related cells. For cooperation, OME promotes shared adapted traits and cellular resources among siblings. For instance, OME was initially discovered for its ability to restore motility to a subset of gliding motility mutants through extracellular complementation, where wild-type (WT) motility proteins from donor cells are transferred to mutants lacking these proteins [3, 4]. OME can also rejuvenate damaged cells with compromised membranes and can transfer adaptation phenotypes by exchanging beneficial components [5, 6]. Conversely, OME also mediates antagonism by exchanging lipoprotein toxins between nonclonal cells that lack a cognate suite of immunity proteins, which are not transferred [7–9].

The sharing of cellular content by OME is an intimate process mediated by the cell surface receptor TraA, which determines recognition specificity. This bidirectional exchange requires direct contact between cells expressing identical or nearly identical TraA receptors. TraA is highly polymorphic, and its variable domain (VD) specifies recognition by homotypic binding [10]. TraA functions alongside its operonic partner, TraB, which is essential for OME but does not influence specificity [11]. Because lipids are exchanged along with proteins, OME is thought to occur by transient OM fusion, with TraAB serving as fusogens [12]. Among the many different proteins transferred by OME are a suite of polymorphic lipoprotein toxins. There are six discrete SitA toxin families, with each family typically represented multiple times in genomes alongside their cognate SitI immunity proteins. Altogether, strains frequent contain over 30 *sitAI* loci and up to 83, resulting in an astronomical degree of discrimination between strains expressing compatible TraA receptors [9]. This interplay between OME and toxin transfer likely drives the evolutionary selection of TraA polymorphisms and recognition specificity.

Myxobacteria form distinct recognition groups based on their TraA receptors. Swapping the VD between *traA* alleles can reprogram recognition specificity. Furthermore, even substituting a single amino acid residue−A/P205−within the VD can alter recognition specificity [13]. To date, experimental studies identified 10 TraA recognition groups, and sequence analysis suggest that many more exist in nature [11]. Despite TraA’s key role in the social interactions of myxobacteria, the molecular basis of recognition specificity remains unclear.

Protein-protein interaction specificity is primarily determined by a subset of residues that strongly covary [14]. These critical residues are typically located at the interface surface between interacting proteins [15, 16]. In this study, we investigated the molecular basis of TraA specificity. To do so, we used AlphaFold, sequence alignments and our experimental findings, to identify candidate residues involved in TraA−TraA recognition. To systematically explore the role of these residues, we created a large, targeted combinatorial library of TraA variants and screened them for altered specificities. Our results reveal that changing a specific subset of residues within the VD is sufficient to reprogram specificity. Interestingly, we found that many of these variants exhibited promiscuous binding properties, capable of recognizing multiple TraA groups. However, most of them could not recognize self, instead switching from homotypic to heterotypic recognition− suggesting that the emergence of novel homotypic recognition is rare. These findings further imply that evolved TraA receptors likely involves intermediate promiscuous states. These intermediates are then selected against, as SitA toxin exchange penalizes heterotypic and/or promiscuous *traA* alleles. Collectively, this work advances our understanding of the molecular and evolutionary basis of social recognition and diversity within

## Results

### Identification of specificity-determining residues

TraA contains a VD, cysteine-rich repeats and a C-terminal protein sorting tag called MYXO-CTERM (Fig. 1A) [3]. We previously showed that TraA−TraA recognition is highly selective, with specificity determined by polymorphisms in the VD [10]. To understand the molecular basis of recognition, we used AlphaFold2 to predict surface residues involved in non-covalent interactions [17]. AlphaFold predicted a TraA−TraA homodimer in a ‘head-to-head’ configuration, formed by approximately a 180° rotation between monomers. The interface residues reside in the VD, and the overall TraA−TraA configuration is analogous to a ‘handshake’ (Fig. 1A, S1). This structure was consistent with our prior analysis that the VD governs cell-cell recognition. By using different alleles of *traA* in AlphaFold predicted structures, we identified 10 surface positions involved in non-covalent TraA−TraA bonds (Fig. 1AB). These interfacial surface residues exhibit considerable sequence variability (Fig. 1B). This analysis suggests that these residues contribute to TraA−TraA recognition specificity, a hypothesis we sought to test experimentally.

**Fig. 1.**
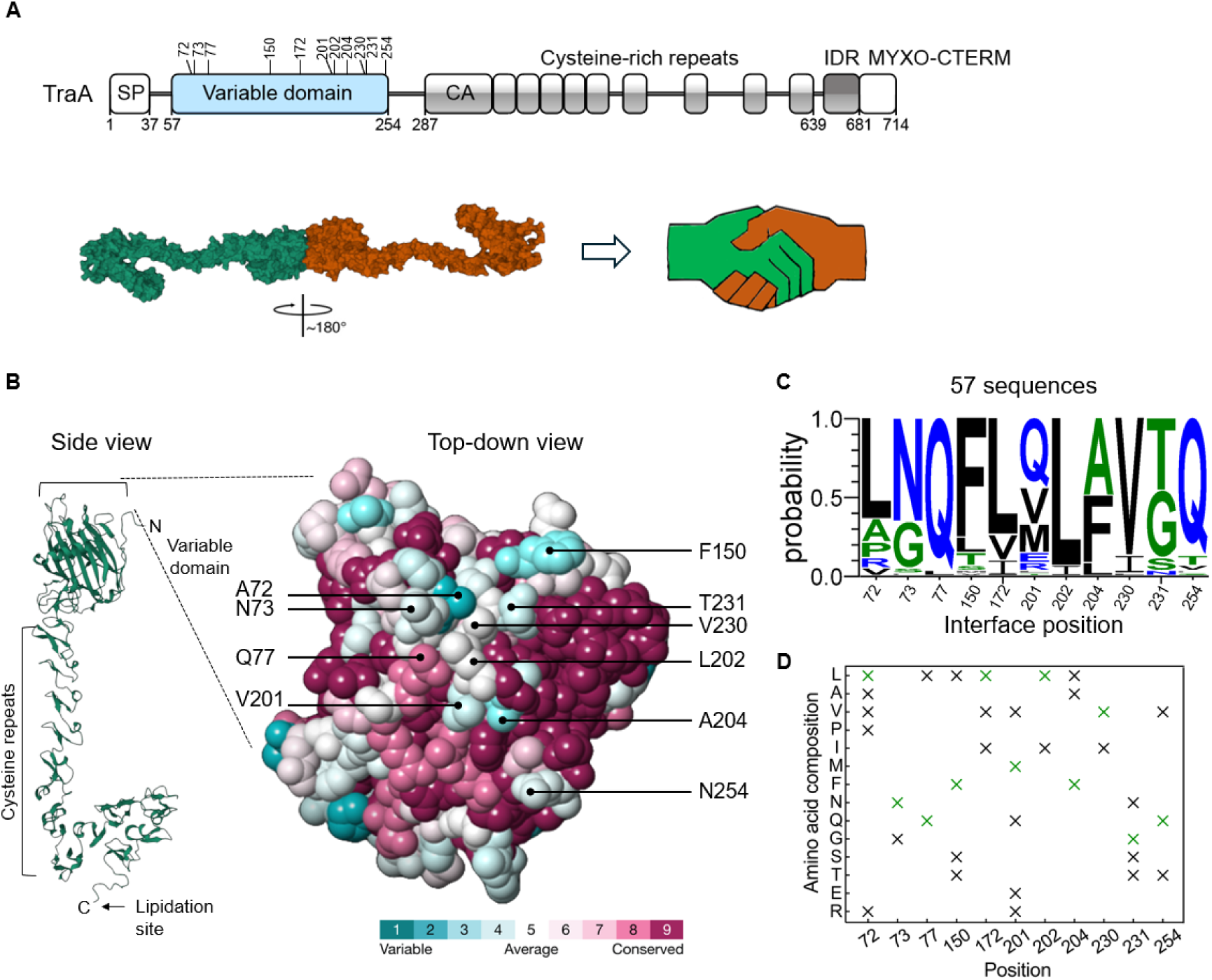
Library rationale. **A**) Domain organization of TraA (group F, *M. fulvus* HW-1 receptor) with residues targeted for substitution in the VD labeled (top), and AlphaFold predicted homodimer TraA structure. IDR, intrinsically disordered region. **B**) Side and top-down view of the VD, where residues degree of conservation are color coded based on a cohort of 57 sequences. **C**) WebLogo of 11 positions targeted for indicated substitutions based on 57 TraA orthologs from group B-F and **D**) shows the 24 substitutions at these positions for library design. Parent sequences in green.

Next, we used sequence alignments to look at the extent of variability among these predicted interface residues. To do so, we restricted the alignment to a cohort of sequences that contained no indels in the VD, which included experimentally determined recognitions groups B−F [13]. This alignment revealed varying degrees of sequence variation at the 10 positions (Fig. S2), as summarized in the WebLogo (Fig. 1C) [18].

### Library construction

Experimentally, we sought to interrogate the role of the 10 positions identified above in recognition specificity. Specifically, we aimed to identify TraA variants that: (i) altered their specificity from one recognition group to another, (ii) were promiscuous, recognizing multiple recognition groups, (iii) engaged in heterotypic binding, failing to recognize self, and (iv) enable a more detailed understanding of the structural basis of specificity to facilitate rational design of new recognition groups.

For library design, we replaced 11 residues from a group F allele (*M. fulvus* HW-1) with residues found in our alignment (Fig. 1C and S2). This strategy limits library complexity by using known amino acids that function at those positions. As an internal control, we included a variable non-interfacial residue, designated as position 5 (aka 172) (Fig. 1C). Each of these 11 positions included 2 to 5 amino acid substitutions in the library (Fig. 1D). Furthermore, in the group F parent allele, we changed the endogenous P205 residue to A205, which abolished recognition to the parent allele (below). This targeted synthetic *traA^F-P205A^* library was constructed by Twist Biosciences, with a theoretical diversity of 172,800 variants, where each clone, on average, contains five substitutions. After amplification and purification from *Escherichia coli*, suicide plasmids were transformed into *M. xanthus* that recombined into the chromosome, generating approximately 32,400 transformants.

### Screen and library assessment

To identify variant *traA* alleles with altered specificity, we employed a ‘stimulation assay’. This assay was based on the ability of donor cells to transfer missing motility proteins to recipient cells by OME, transiently restoring gliding motility. To test the sensitivity in our screen, we mixed donor and recipient strains at various ratios. Both strains were nonmotile and shared the same *traA* allele; where the Δ*cglC* Δ*tgl* recipient was rescued by OME transfer of CglC and Tgl from the donor, restoring A- and S-motility (Fig. S3A). Strikingly, rare but positive recipients were detected at donor:recipient ratios as low as 1 : 200,000 cells (Fig. S3B). We conclude that our assay was highly sensitive and capable of detecting rare positive *traA* clones with altered specificities from large pooled libraries.

To assess library quality and screen feasibility, we randomly selected 50 clones for characterization. Of these, 15 (30%) contained unintended *traA* mutations and were excluded from further analysis (Fig. 2A). Among the remaining clones, 22 (44%) recognized one or more members from a donor pool consisting of TraA A-F recognition groups (Fig. 2A, B). The remaining 12 clones exhibited unique recognition patterns, as they did not recognize groups A-F but instead recognized variant alleles from within the library itself (Fig. 2A, B). These unexpected results indicated our library contained many *traA* alleles with altered or novel specificities.

**Fig. 2.**
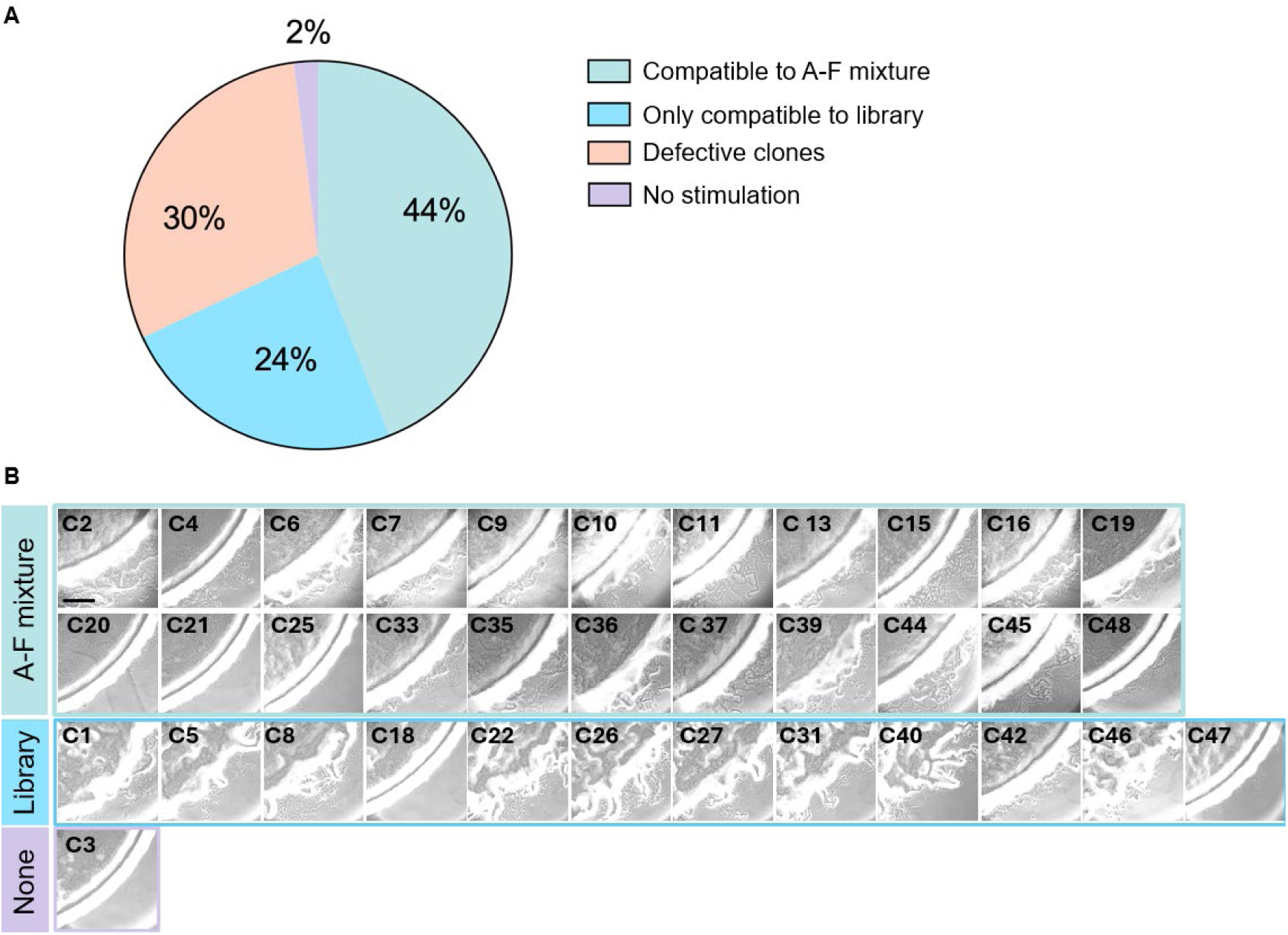
Library characterization by stimulation assays. **A**) Overview of 50 randomly selected clones tested for activity against a pool of recipients belonging to recognition groups A-F or library pool. **B**) Stimulation results for clones from panel A. Scale bar, 200 µm.

### TraA variants with altered recognitions

The *traA^F-P205A^* library was screened against four different TraA groups− A, B, C and E (Fig. 3A, B). Group D was excluded from screening due to low-level crosstalk with the parent library allele *traA^F-P205A^* (Fig. 3C), and the original WT group F was also excluded. All six TraA recognition groups are found in *M. xanthus* isolates. Notably, group A, which originates from the lab strain DK1622, has five indels compared to the other five groups and is phylogenetically more distant (Fig. S4). Across the four library screens, variants with altered recognition specificities were readily identified, even though the parent allele *traA^F-P205A^* did not recognize groups A, B, C or E (Fig. 3C). In total, 62 clones were retained for further characterization.

**Fig. 3.**
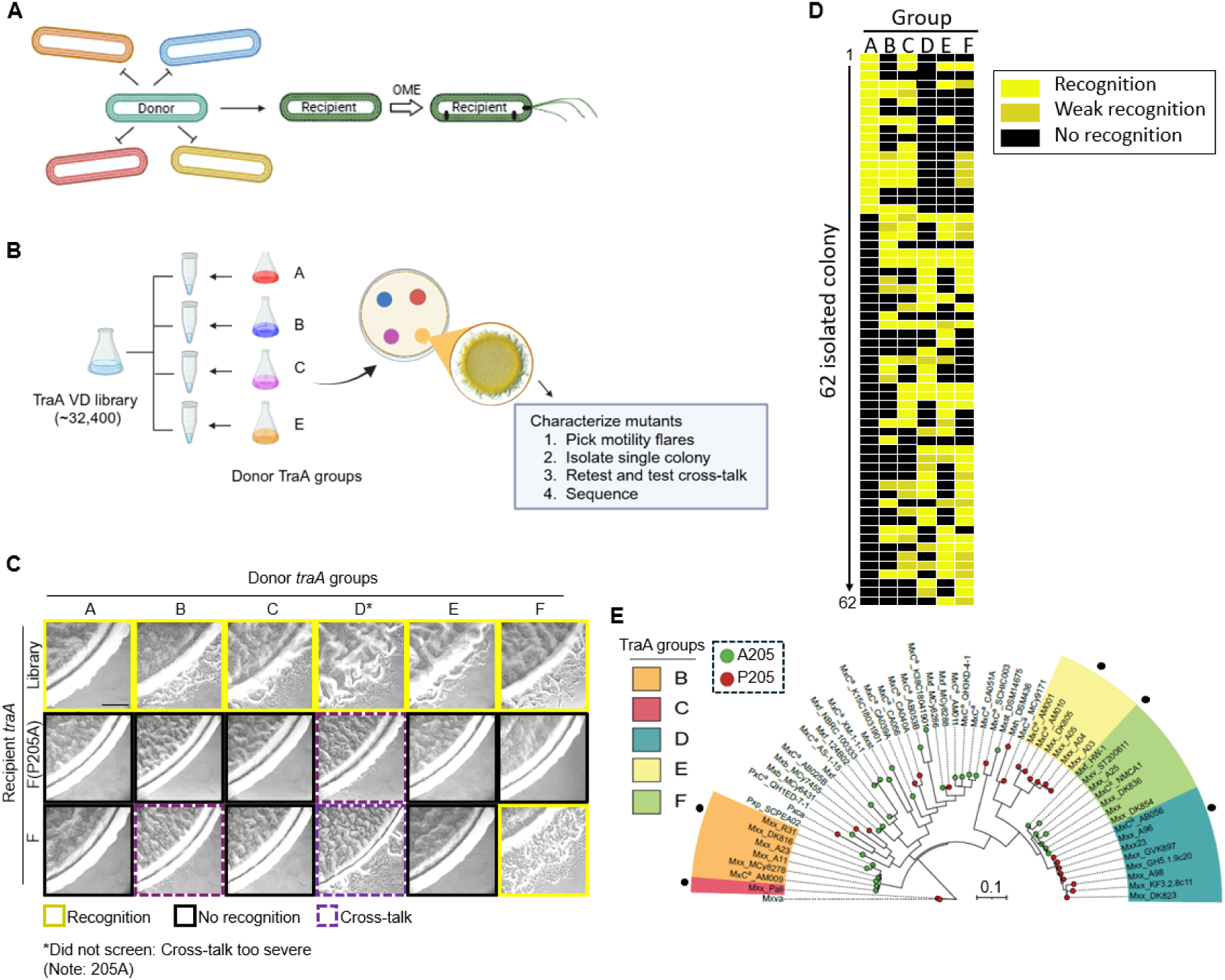
Library screen and recognition profiles. **A**) Schematic of library screen. Only recipients in library with compatible TraA receptor to given donor receptor will engage in OME that rescues motility. **B**) Overview of the screening strategy against TraA recognition groups A, B, C, and E. **C**) Stimulation tests of library compatibility against TraA groups A–F. As controls, the original F receptor and the F^P205A^ substituted receptor backbone used for library construction are shown. Scale bar, 200 µm. **D**) Summary of 62 positive library clones tested against TraA groups A–F. Clones were isolated from screens against groups A, B, C, and E as shown in C. **E**) Maximum likelihood tree based on 57 VDs of *Myxococcaceae* TraA orthologs used for library design (Fig. 1). Each allele because they contain no indels. Experimentally determined recognition groups are color coded. Black dots indicate the corresponding strain used in panels C and D. Scale bar, number of substitutions per residue.

To investigate recognition specificity, these clones were tested against TraA recognition groups A through F in stimulation assays. A summary of the 62 clones’ recognition profiles is shown in Fig. 3D. Additionally, the sequence relationships of these five parent groups among 57 natural alleles is shown in Fig. 3E. This analysis revealed that the majority of TraA variants exhibited promiscuous interactions. In fact, only 7 out of 62 clones showed specificity exclusively to the recognition group from which they were isolated. In contrast, 23 clones recognized four or five different recognition groups. We conclude that the targeted residues selected for substitutions that play key roles in recognition specificity, and that such changes often lead to promiscuous heterotypic binding.

These positive clones were sequenced, and the amino acid biases at each of the 11 positions were visualized with WebLogos (Fig. 4A). Compared to the initial library composition, several positions showed greater than 2-fold enrichments in groups A, B, and C. For instance, group A-positive clones consistently contained a serine at position 10 in all 18 clones. In contrast, groups D, E and F exhibited negligible preferences compared to the original library composition. At the control residue, there was no significant change at position 5 in either the amino acid composition or fold change. From this analysis, we identified positions and amino acids that were important for recognition specificity. Additionally, residues absent from WebLogos were inferred to have a detrimental effect on specificity in those recognition groups.

**Fig. 4.**
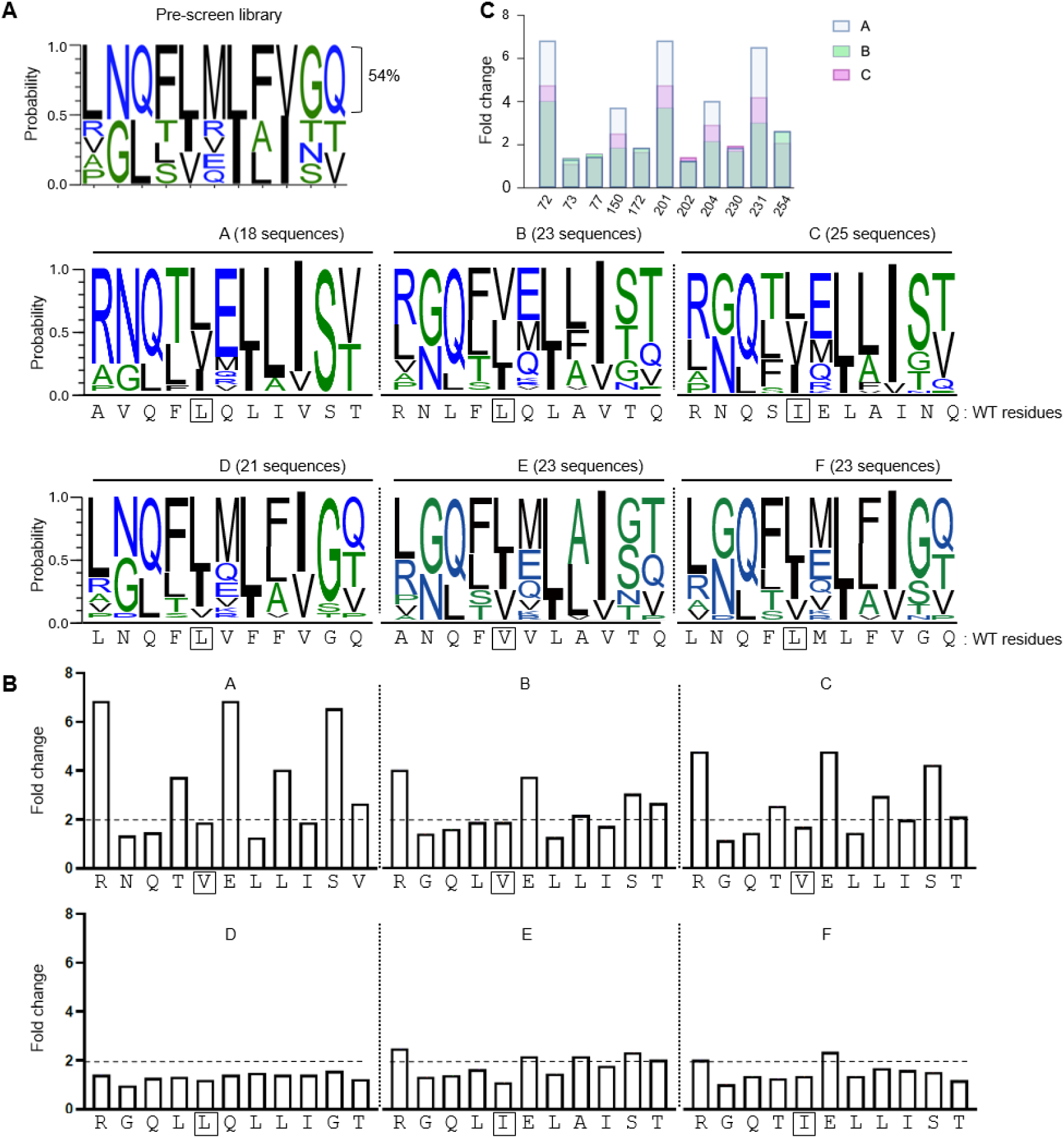
Enrichment of amino acids for the different TraA recognition groups. **A**) WebLogos of amino acid preferences from TraA positive clones against groups shown. **B**) Amino acid enrichment and fold-change highlighted from TraA-positive clones for each group. Amino acids with the highest fold-change enrichment at each position are shown (bottom of graphs). Control position boxed. **C**) Fold-change highlighting amino acid enrichment in group A, B and C.

To determine which positions were important for recognition, the fold change of preferred amino acids was calculated (Fig. 4B). Significant enrichments were observed at same six positions in groups A and C, five in group B, and no significant changes were found in groups D, E and F. These findings correlated with the phylogenetic relationships or distance among these six parent alleles (Fig. S4B). Additionally, crosstalk among receptors was expected to be more likely among closely related proteins [19], a trend we generally found in the screening results (Fig. 3D). Finally, since receptors from groups A, C and B were more distantly related to the F^P205A^ based library than receptors from groups D, E and F, the former exhibited a greater degree of amino acid substitutions biases than the latter.

### Homotypic versus heterotypic allele recognition

In nature, recognition occurs by homotypic binding by identical TraA receptors, which helps to ensure cells are clonal. In contrast, our screen readily identified alleles involved in heterotypic allele recognition. To test for self (homotypic) recognition, we selected 12 of the 50 random isolates that were incompatible with groups A−F but nevertheless were functional when screened against the library (Fig. 2). Interestingly, none of these 12 variants recognized self, but nine exhibited heterotypic recognition, with eight being promiscuous (Fig. 5A). Next, we selected another 12 clones from the pool of 62 identified from screening against group A, B, C or E (Fig. 3D). Among these 12, five recognized only one group, while the other seven were promiscuous. This analysis revealed that 4 of 7 of the promiscuous clones recognized self to varying degrees, while only 1 of the 5 selective clones exhibited weak self-recognition (Fig. 5B, C). We conclude heterotypic allele interaction were favored over homotypic recognition when specificity residues were changed, although promiscuous clones were more likely to also self-recognize.

**Fig. 5.**
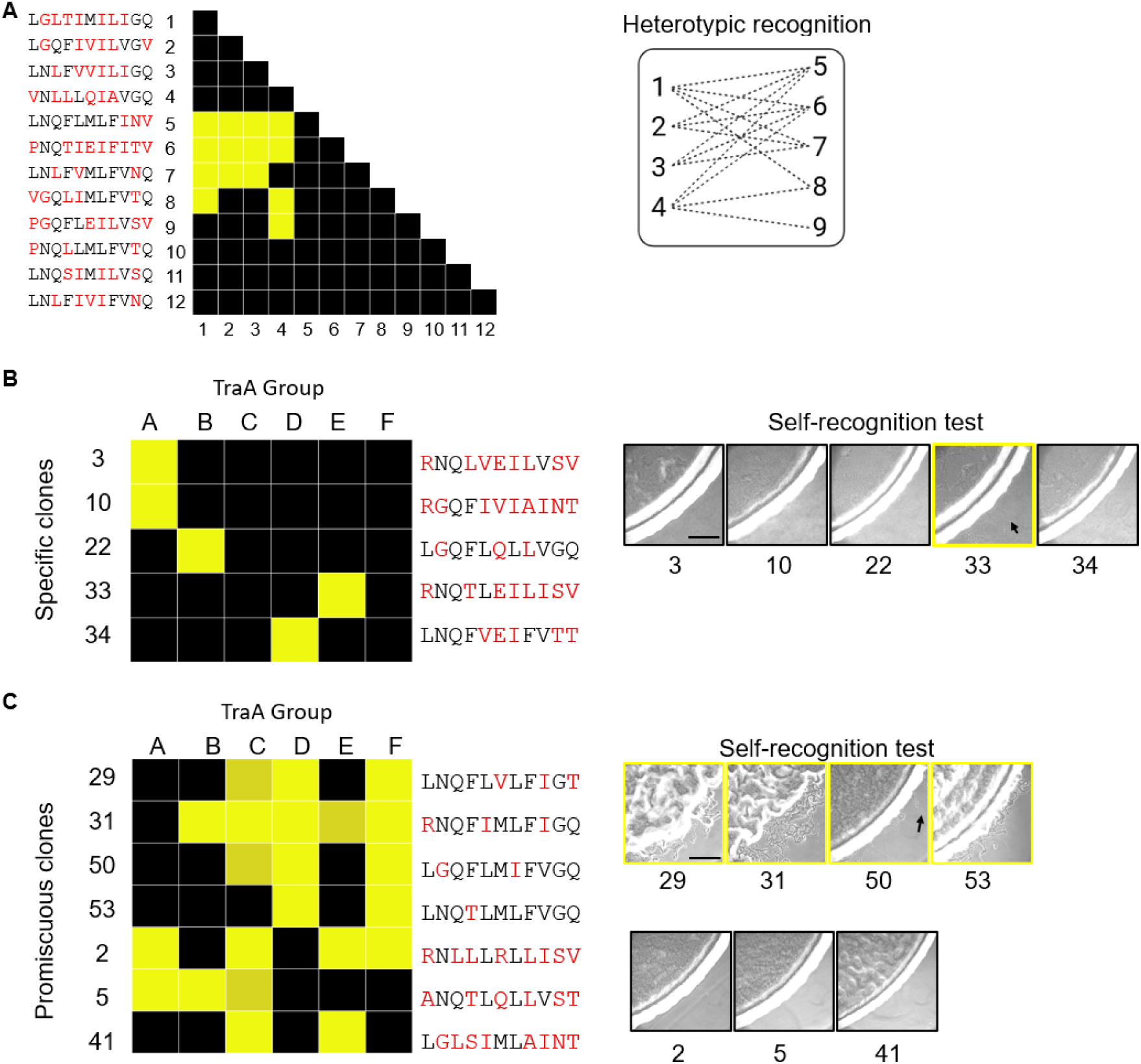
Ability of TraA variants to recognize self or homotypic binding. **A**) Twelve functional variants against the library, but not active against groups A-F (from Fig. 2). **B**) TraA variants that only recognize one group among A-F (numbers refer rows/clones from Fig. 3D) were tested for homotypic recognition (right panel). **C**) TraA promiscuous variants (numbers refer rows/clones from Fig. 3D) tested for homotypic recognition (right panels). Tables: yellow, stimulation; shaded yellow, poor stimulation; black, no stimulation. Red letters, substituted amino acids. Scale bar, 200 µm.

### Distant TraA recognition groups were not library compatible

Given the ease of identifying library variants that recognize groups A through F, we sought to expand the screen against distant TraA recognition groups G through J, which originated from genera outside of *Myxococcus* (Fig. 6A) [20]. Using the same sensitive stimulation assay, we were unable to identify positive clones from our library (∼32,400 variants) against these four divergent TraA receptors (Fig. 6B). We conclude, that these alleles, which contain various numbers of indels (Fig. S4), were too divergent to interact with our *traA^F-P205A^* based library.

**Fig. 6.**
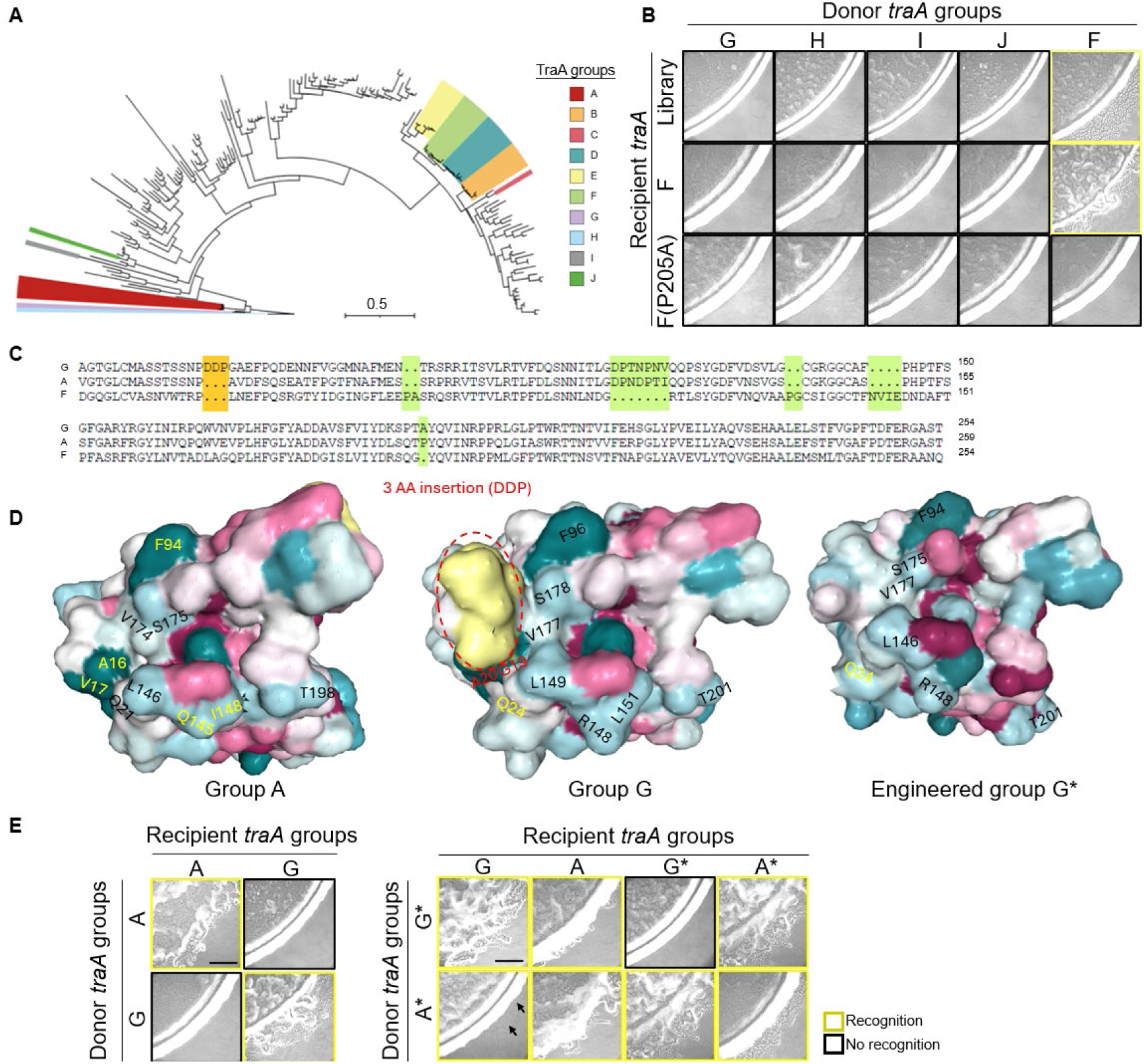
Rational design of TraA recognition alleles. **A**) Maximum likelihood tree of the VDs from 193 *Myxococcaceae* TraA orthologs that includes sequences with indels. TraA groups A–J color-coded. Scale bar represents the number of substitutions per amino acid site. **B**) Library screen against TraA groups G-J. **C**) VD sequence alignment highlighting variations and indels between groups A, F and G. **D**) AlphaFold predicted structural differences in the VD region among TraA groups A, G and G*. **E**) Recognition outcomes of engineered variants TraA-A*, TraA-G* and their parent alleles based on stimulation assays. Black arrows, stimulated flares; scale bars, 200 µm.

### Engineered TraA receptor

It was noteworthy that we readily isolated variants that interacted with TraA recognition group A, which contains five indels in the VD compared to the library and was phylogenetically distant from groups B-F (Fig. 6A), but we were unable to identify positive clones from groups G through J, which had a comparable number of indels (Fig. S4). This raised the question: how did the library recognized group A, but not these other groups?

To investigate the molecular basis of library incompatibility, we focused on sequence and structural differences between group A and G. Representative alleles from these two groups were similar in that they both shared five indels when compared to groups B-F (Fig. 6C and S4). When comparing only the A and G sequences, there was a single three-amino acid indel between them (Fig. 6C). In AlphaFold structural predictions, these three residues formed a striking bulge on the predicted interface surface (Fig. 6D). Given our ability to identify library variants that recognize group A, and the presence of three amino acid insertions as an obvious sequence difference between group A and G, we hypothesized these residues played a key discriminating role preventing promiscuous TraA−TraA interactions. To test this idea, we constructed an in-frame deletion that removed these amino acids (D69, D70, and P71; Fig. 6C) and substituted the two adjacent interface residues with those found in the *traA^F-P205A^* library (DDPGA → LN). Strikingly, this mutant (G*) displayed a relaxed specificity toward group A, and it retained its ability to recognize the parent group G receptor (Fig. 6E). However, the G* mutant did not recognize self, thus it transitioned from homotypic to heterotypic allele recognition. We additionally constructed a reciprocal allele in group A (adding DDP before AV, called A*), which showed homotypic and heterotypic recognition with A*, A, G and G*. Thus, these variants exhibited promiscuous interactions compared to their WT alleles.

### Cell–cell adhesion and visualization of OME

TraA homotypic recognition promotes cell-cell adhesion [13, 20]. Here we tested whether heterotypic recognition also forms cell-cell adhesions. The *traA^3^* variant strain (Fig. 5B) was cultured or cocultured with a *traA^F-P205A^* strain labeled with mCherry. We found that *traA^3^* cells did not bind to themselves, nor to the parent *traA^F-P205A^* strain (Fig. 7B). However, the *traA^3^* strain showed heterotypic binding when cocultured with a *traA^A^* strain labeled with GFP, as implied by stimulation results (Figs. 3D and 5B). These findings contrast with homotypic binding, where cell adhesion occurs only between identical TraA receptors (Fig. 7A) [13].

**Fig. 7.**
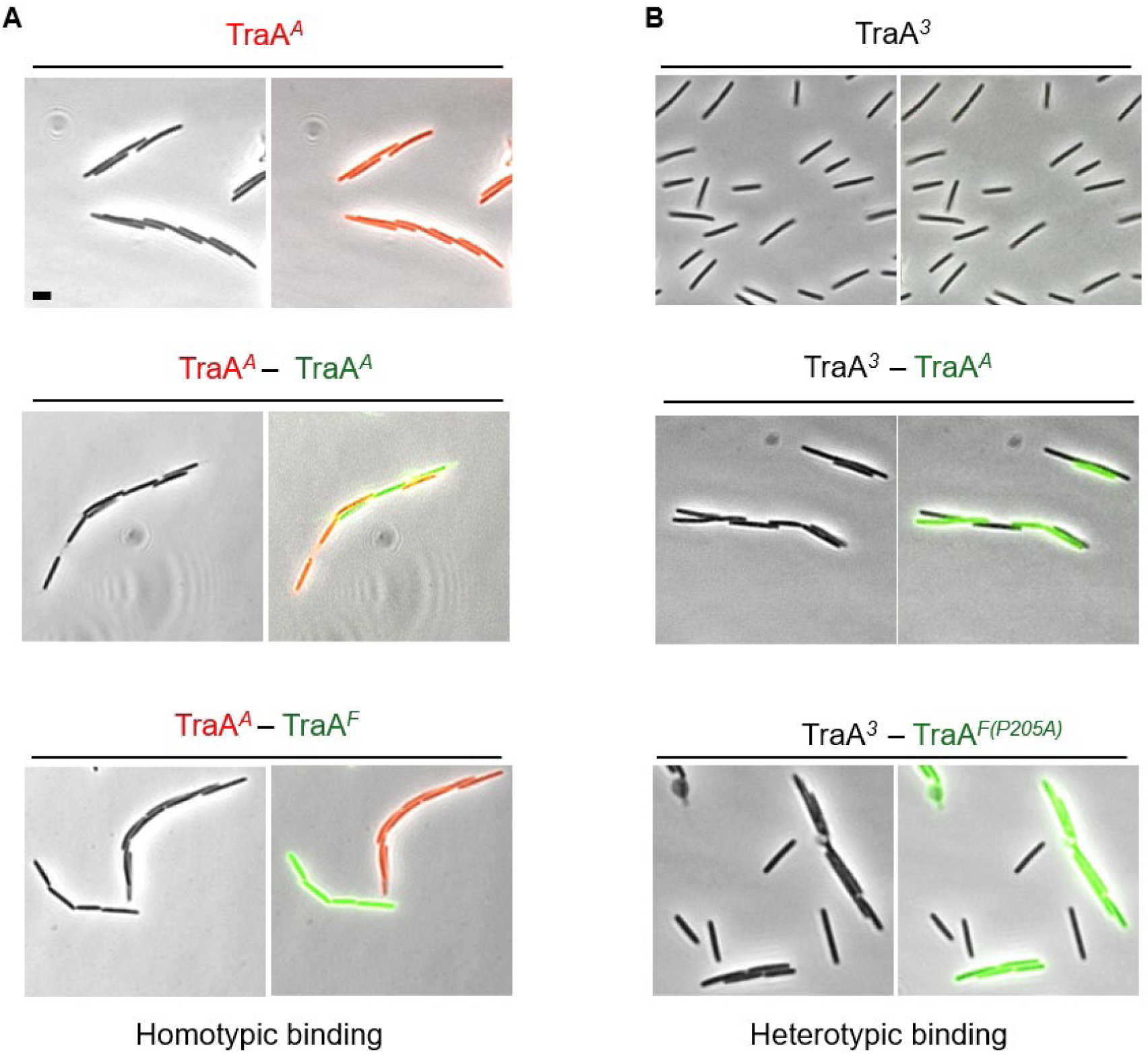
Homotypic to heterotypic switch for cell adhesion. **A**) *traA^A^* strain (mCherry) cultured alone, cocultured with *traA^A^* (sfGFP), or with *traA^F^* strain (sfGFP). Superscript designates recognition group. **B**) A library variant *traA^3^* cultured alone, with *traA^A^* (mCherry), or with *traA^F-P205A^* (sfGFP). Scale bar = 2 µm.

TraA recognition leads to the exchange of OM goods [20]. Here, we sought to visualize heterotypic recognition and OME between strains. To do so, we labeled populations with SS_OM_-GFP or SS_OM_-mCherry lipoprotein reporters that are readily transferable. Initially, following mixing and plating, cells were phenotypically distinct (red or green). Strikingly, *traA^A^* and *traA^3^* cells became homogeneous and phenotypically indistinguishable after 90 min (Fig. 8B). In contrast, no OME occurred between cells expressing *traA^3^* and *traA^F-P205A^*. As controls, OME occurred between cells expressing same the TraA receptors but not between cells expressing different alleles (Fig. 8A). Together, these findings confirm TraA variants can switched binding partners from homotypic to heterotypic.

**Fig. 8.**
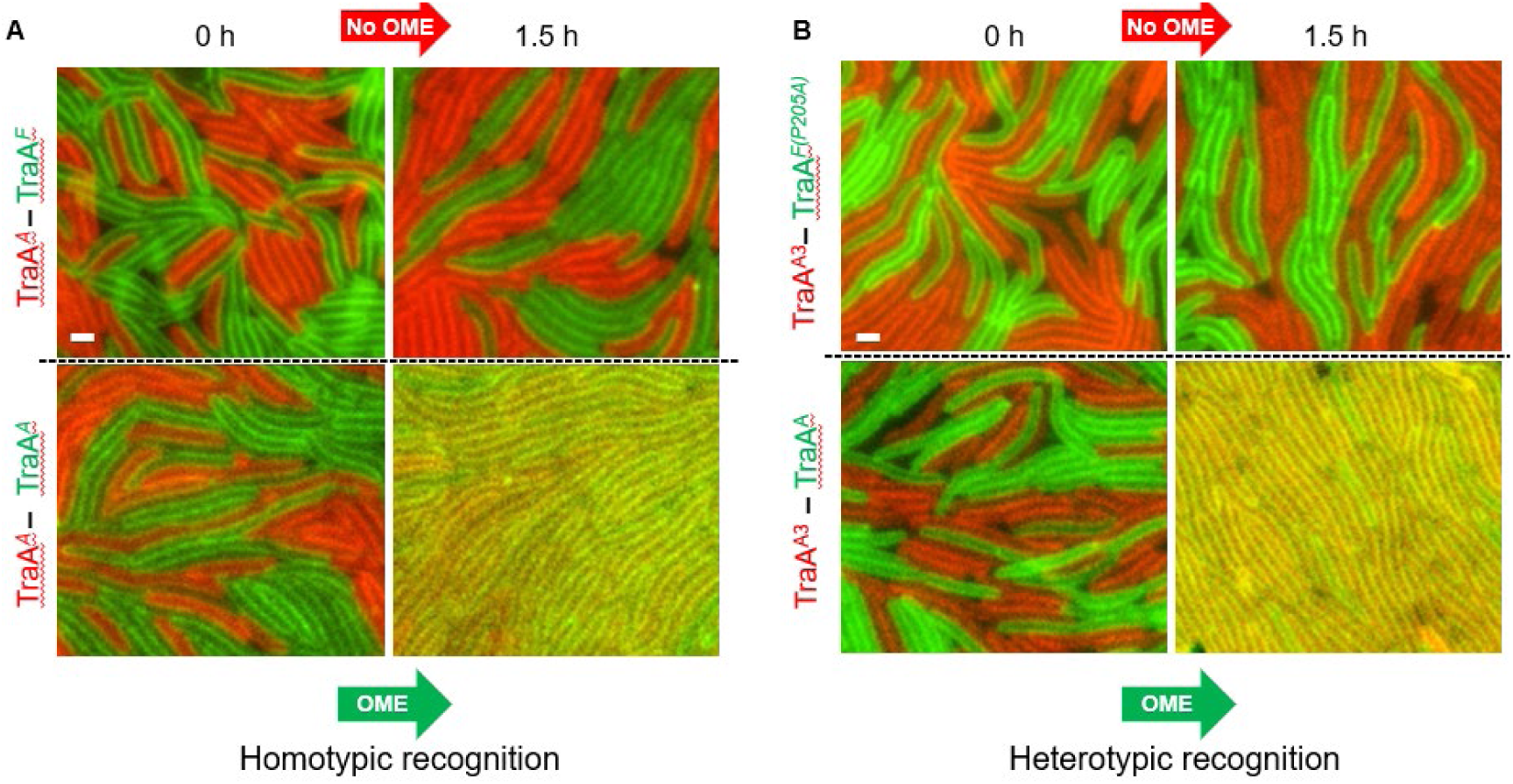
Homotypic versus heterotypic OME between kin populations harboring transferable SS_OM_-GFP or SS_OM_-mCherry reporters. **A**) TraA homotypic recognition governs OME. *traA^A^* (mCherry) mixed with *traA^F^* (sfGFP), or with *traA^A^* (sfGFP). **B**) TraA heterotypic recognition governs OME. A library variant *traA^3^* (mCherry) mixed with *traA^F-P205A^* (sfGFP), or with *traA^A^* (sfGFP). Incompatible TraA receptors block exchange after the indicated time (top panels), whereas compatible receptors transform into a homogeneous tissue-like population through OME (orange) (bottom right panels). See Supplementary Fig. 5 and 6 for single channel micrographs and Supplementary Table 1 for strain details. Scale bar = 1 µm.

## Discussion

The specificity of cell-cell recognition for OME in *M. xanthus* is mediated by the cell surface receptor TraA, which enables neighboring cells to distinguish self from nonself. This recognition, in turn, leads to benefits or harmful interactions directed towards kin. Consequently, *traA* is often described as a ‘greenbeard’ social gene [21, 22]. Consistent with this role, TraA is polymorphic, and the protein is amendable to substitutions and indels, which has allowed it to evolve numerous recognition specificities. Previous mutagenesis studies showed that specificity can be changed by swapping VD domains, altering residues, or even with single amino acid substitutions at a key recognition switch residue [13]. However, the molecular basis of dozens − if not hundreds−of different specific homotypic recognition groups remained largely unknown. In this study, we used sequence alignments and AlphaFold-based modeling to predict amino acid residues involved in TraA-TraA recognition. We then designed a combinatorial library to survey and test these residues in recognition and specificity changes. Notably, we found that sequence variations in TraA frequently generate promiscuous variants that interact with multiple recognition groups. However, in nature, promiscuous TraA receptors are not observed, indicating there are selective pressures that favor a dynamic process for the evolution of specific homotypic binding.

The evolution of protein interfaces is a tightly constrained process, particularly when interfaces contribute not only to molecular binding but also to broader cellular functions [23]. These constraints are often reflected in lower mutation rates observed at interface regions compared to other regions, highlighting the need for coordinated coevolution with interaction partners [24]. In myxobacteria, the evolution of TraA is subject to similar constraints, as changes in its interface residues influence cell cooperation and antagonism.

Based on our findings, we offer a rationale for the natural evolution of TraA polymorphisms while maintaining homotypic specificity−a contrast to our experimental observations (Fig. 9A). We suggest that during TraA evolution, sequence variants fall into four categories: homotypic selective, heterotypic selective, heterotypic promiscuous, and homotypic promiscuous. Homotypic selective represents the WT state, which enhances population fitness by promoting cooperation among siblings while remaining neutral toward non-siblings with divergent TraA receptors. Homotypic promiscuous variants support sibling cooperation but may cause harm or antagonism with non-siblings due to OME of polymorphic SitA toxins. Heterotypic selective is limited to specific strains with a different *traA* allele, fostering OME-mediated antagonism. In contrast, heterotypic promiscuous recognition is the most detrimental, as TraA fails to recognize self but interacts with and antagonizes strains with different TraA receptors. We propose that these promiscuous and heterotypic interactions−frequently encountered in our screen−represent intermediate states in the evolution toward homotypic specificity, which occurs less frequently.

**Fig. 9.**
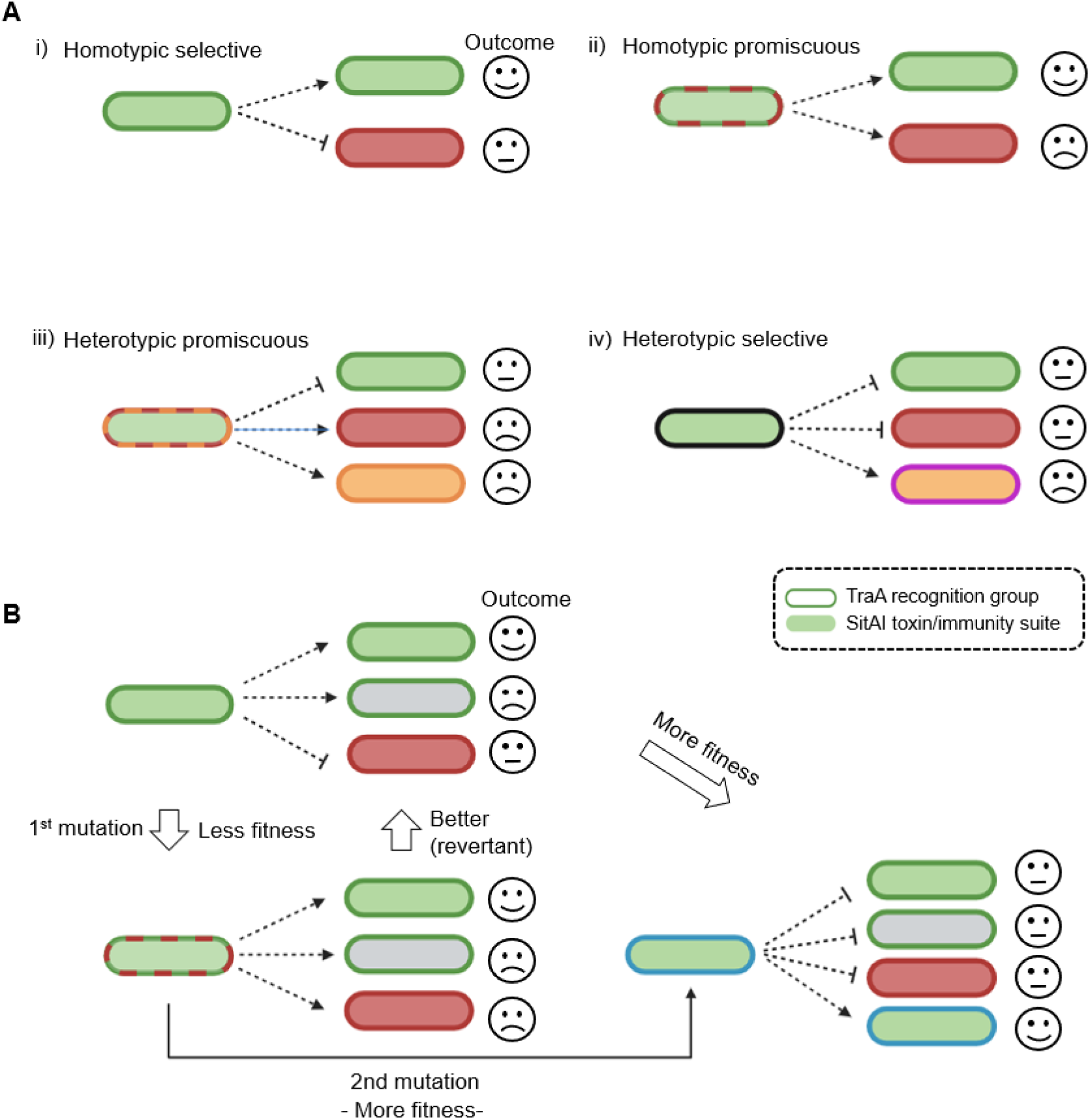
Evolutionary scenarios and selective forces governing TraA homotypic recognition. **A**) Four types of TraA recognition during evolutionary diversification. **B**) Model illustrating the evolutionary steps to create a new homotypic TraA– TraA recognition specificity through promiscuous intermediate. See text for details.

We propose that new TraA specificities typically evolve through the generation of promiscuous variants, driven by a reward-and-punishment system arising from cell-cell interactions. These evolutionary steps ultimately enhance overall population fitness (Fig. 9B). In wild myxobacteria populations, homotypic selective recognition produces three outcomes; promoting cooperation among siblings, antagonizing individuals with compatible TraA receptors but different SitA toxins, and maintaining neutrality toward non-siblings with divergent TraA groups. TraA diversification may proceed through a series of evolutionary changes, where the initial mutation broadens recognition among a wider range of TraA variants. However, this promiscuity increases the frequency of antagonism due to a diverse set SitA toxin repertoires. Therefore, subsequent *traA* mutation(s) are selected that narrows specificity back toward self, resulting in novel homotypic recognition that promotes sibling cooperation while remaining neutral toward others (Fig. 9B).

Our findings challenge existing models for the evolution of protein-protein interactions [16, 25, 26]. In coevolution models involving compensatory mutations, one protein can lose its function until a corresponding mutation in its partner restores the interaction and hence, function [25, 27]. In contrast, the promiscuous intermediate model involves changes in protein specificity without passing through a non-functional state [16]. This model is exemplified in toxin-antitoxin systems, where interaction specificity evolves through promiscuous intermediates because retaining the interaction is essential for cell viability. Our results suggest that for TraA recognition, compensatory evolution can also occur where self-recognition is transiently lost or reduced while promiscuity emerges. Such promiscuous alleles likely represents short-lived evolutionary advantages that prevent harmful homotypic interactions with aggressor cells [28], including from ancestor cells that recently acquired a new *sitAI* locus via horizontal gene transfer [8, 28]. However, this promiscuity is unlikely to persist, as OME interactions with other strains lead to harmful non-sibling engagements. This, in turn, creates selective pressures to restore homotypic self-recognition, potentially resulting in a new specificity group. The promiscuous variants described here differ from a previously described specificity switch at the conserved A/P205 position, which occurs via a single-step mutation [13].

Our results demonstrate that analyzing amino acid variations from large multiple sequence alignments and AlphaFold structure predictions can effectively guide the identification of specificity determinants in protein-protein interactions. This approach enabled us to identify a cluster of 10 residues particular, we found that six positions primarily dictated specificity of groups A, B and C. In another system, interaction specificity in one family was narrowed down to five residues in the toxin and four in the antitoxin, which can mutate and adapt to each other without compromising protein function [16]. Moreover, partner specificity in paralogous families of bacterial histidine kinase-response regulator complexes is determined by just three residues in the complex interfaces [29].

Insertions and deletions represent another evolutionary mechanism that contributes to the development of new specificities [23, 30, 31]. Here, we uncovered an intriguing example in which a TraA^G^ group member contained a three-amino-acid insertion that resulted in a bulge on the recognition interface. This bulge appears to sterically hinder binding to other recognition groups and thus provides a structural basis for specificity. We tested this with an engineered TraA^G*^ deletion variant (DDPGA → LN) and showed that it became promiscuous−recognizing group A receptors while still weakly recognizing its parent allele. These results suggest that the ‘bulge’ residues indeed play a key role in excluding nonspecific or heterotypic binding. Our findings open the door for future rational engineering of new TraA recognition receptors and possibly other families of interacting proteins.

## Materials and Methods

### Bacterial strains and growth conditions

Table S1 lists bacterial strains and plasmids used in this study. *M. xanthus* strains were cultured in the dark at 33 °C with shaking in CTT media (1% [w/v] casitone, 1 mM KH_2_PO_4_, 8 mM MgSO_4_, 10 mM Tris-HCl pH 7.6). For ½ CTT CTT, casitone was reduced to 0.5%. *E. coli* cultures grown in Luria broth (LB) medium at 37 °C for cloning. For solid media plates, agar was added at 1.5% or 0.5% (wt/vol). Antibiotics were added at the following concentrations when indicated: 50 μg/ml kanamycin (Km) for *M. xanthus* and *E. coli*, 15 ug/ml oxytetracycline (oTc) for *M. xanthus*, and 10 ug/ml tetracycline (Tc) for *E. coli.* TPM buffer (CTT without casitone) was used to wash cells.

### Plasmid and strain construction

Primers used in this study are given in Table S2. The *traA^F-P205A^* library was cloned onto pMR3487 [32] with the *traB* gene and expression was driven by an inducible P_IPTG_ promoter. To create plasmid backbone for mutant library, the fragment of *traAB^Mf^* from pPC4 was amplified by PCR. The resulting fragment was then sub-cloned into pMR3487 (linearized with XbaI and KpnI) in Gibson Assembly Master Mix (New England Biolabs).

The TraA-G* (DDPGA → LN) variant was generated by deleting codons 69-73 from plasmid pPC26 and inserting the codons CTC and AAC coding for leucine and asparagine, respectively. Primers with substituted bases were used for PCR amplification. The resulting fragments were assembled using Gibson Assembly Master Mix.

To generate TraA-A* (insertion of DDP before AV) plasmid, primers used for PCR amplification were engineered with codons GAT, GAC and CCG coding for aspartic acid, aspartic acid and proline, respectively. The resulting fragments were assembled via Gibson Assembly.

Markerless in-frame deletion of *cglC* was constructed in DK1253 strain (*pilQ1*, *sglA1*, *tgl1*). Briefly, pΔ*cglC*, containing a *cglC* deletion cassette and a Km^R^-galK selection cassette, was electroporated into DK1253. Homologous recombinants were first selected by Km^R^, and then plasmid excision was counter-selected on 3% galactose plates. This markerless deletion of *cglC* generated DW2229, which was confirmed by PCR with flanking primers and subsequent phenotype analysis. To create markerless in-frame deletion of *traA* in DW2229 (*pilQ1*, *sglA1*, *tgl1*, Δ*cglC*), pDP28 containing *traA* deletion cassette was transformed in DW2229, and selected in a similar manner as described above.

All plasmids were verified by PCR, restriction enzyme digestion, and DNA sequencing. Verified plasmids were then electroporated into *M. xanthus* cells and selected with appropriate antibiotics.

### Phylogenetic analysis

Maximum likelihood (ML) phylogenetic trees were constructed in PhyML 3.0 [33]. We first identified the best-fit models based on the Akaike Information Criterion (AIC) in ProTest 3 [34]. For the analysis of 57 TraA VD orthologs from the *Myxococcaceae* family (Fig. 3A), the best fitting model (Q.insect+R+F) selected by ProtTest was used, and 1,000 bootstrap replicates were performed. In addition, sequences of the 193 TraA VDs spanning of *Cystobacterineae* suborder were used to generate a ML tree. For this analysis, the model (Q.yeast+R+F) was selected, and 1,000 bootstrap replicates were performed. Phylogenetic trees were visualized with iTOL [35].

### Stimulation assay (also for screening conditions)

This assay was essentially performed as described [11]. Briefly, *M. xanthus* was grown overnight to mid-log phase. Nonmotile non-stimulatable donor strains were mixed with nonmotile stimulatable recipient strains at a 1:1 ratio, and placed onto ½ CTT agar plates containing 2 mM CaCl_2_. After incubation at 33 °C, the edges of colonies were imaged with Nikon E800 phase contrast microscope with a 10× objective lens coupled to a digital imaging system.

### Cell-cell adhesion assay

Strains and cocultures were mixed as indicated and incubated overnight with shaking at 300 rpm at 33 °C. Cultures were grown to mid-log phase at a density of 1 × 10^7^ cells per mL. Cell suspensions were directly mounted on glass slides and examined using a Nikon E800 microscope equipped with a 60× oil objective lens and FITC or Texas Red filter sets coupled to an imaging system.

### Protein transfer assay

Transfer assays done as described [20]. In brief, to test SS_OM_-mCherry or SS_OM_-GFP transfer, strains were grown to mid-log phase (∼5 × 10^8^ cells per ml) and mixed as indicated and spotted on ½ CTT 1.2% agar pads. Images taken after spots dried on the agar pads, or incubated at 33 °C for indicated times.

## Acknowledgments

We thank Osmond Ekwebelem for structural modeling assistance and Pengbo Cao and Wall lab members for helpful suggestions. This work was supported by the National Institutes of Health grant GM140886 to D.W.

## Author contributions

T.G. and D.W. designed experiments, analyzed data, and wrote the paper. T.G. performed experiments.

