## Supplemental material for "Experimental evolution of promiscuous kin recognition from a homotypic specific cell surface receptor"

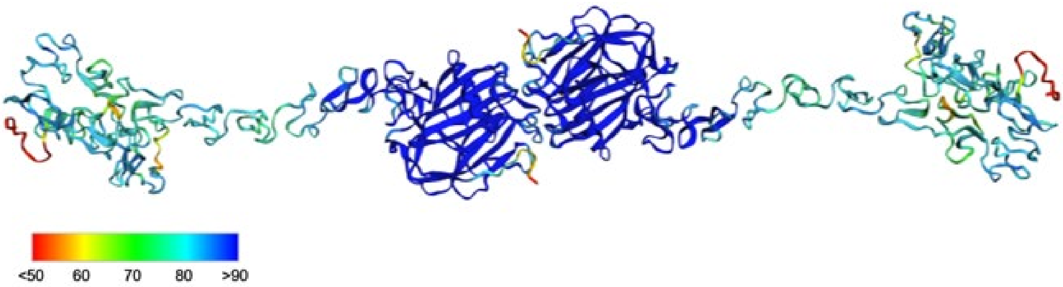


**Fig. S1** Backbone representation of TraA-TraA homotypic binding. Structure predicted by AlphaFold2 and color-coded by pLDDT confidence scores.

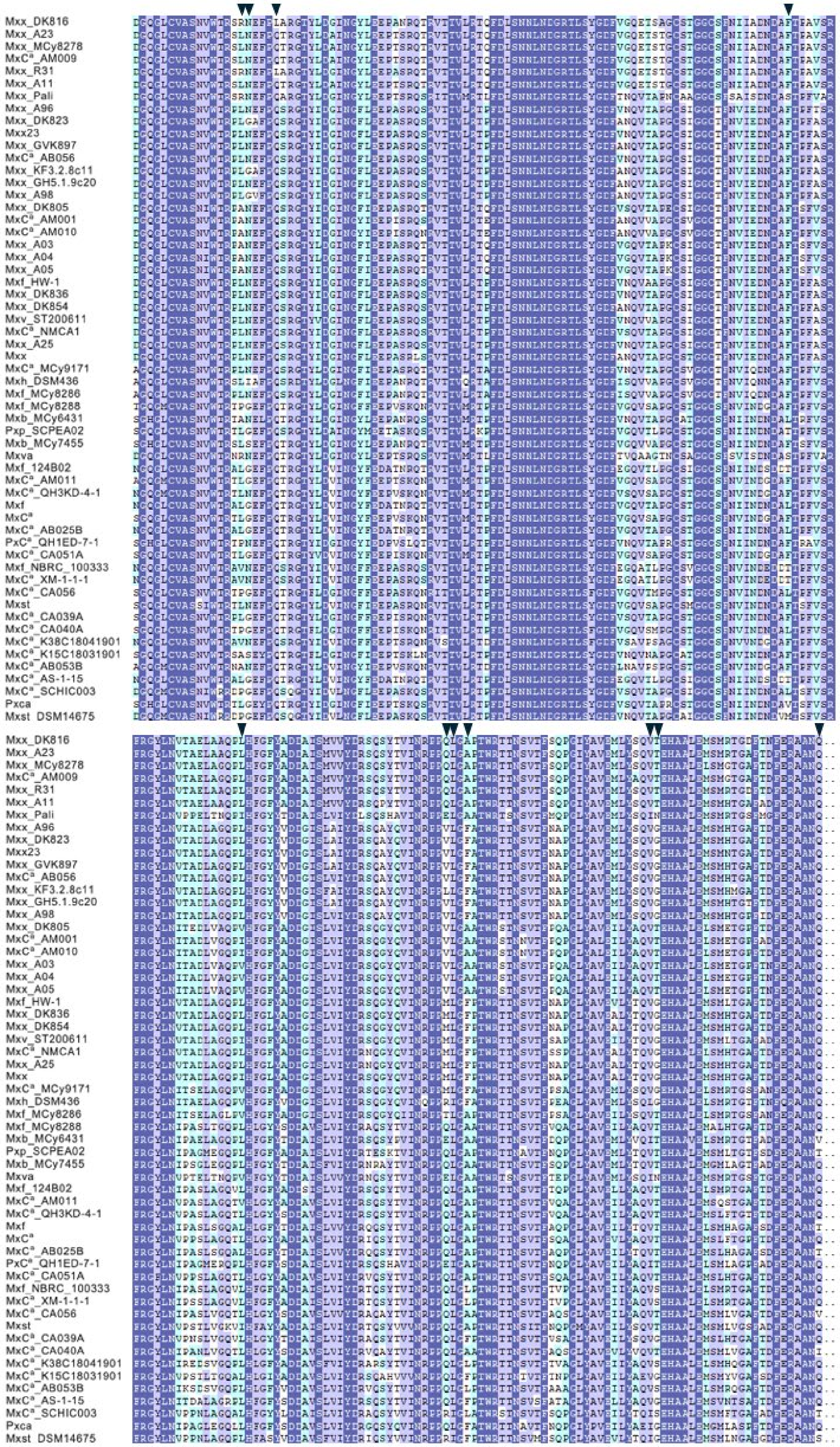


**Fig. S2** Sequence alignment of the TraA VD from groups B to F across 57 sequences that contain no indels. Black triangles, 11 positions substituted for library construction. Homologies: Blue, 100%; violet, >75%; cyan >33%.

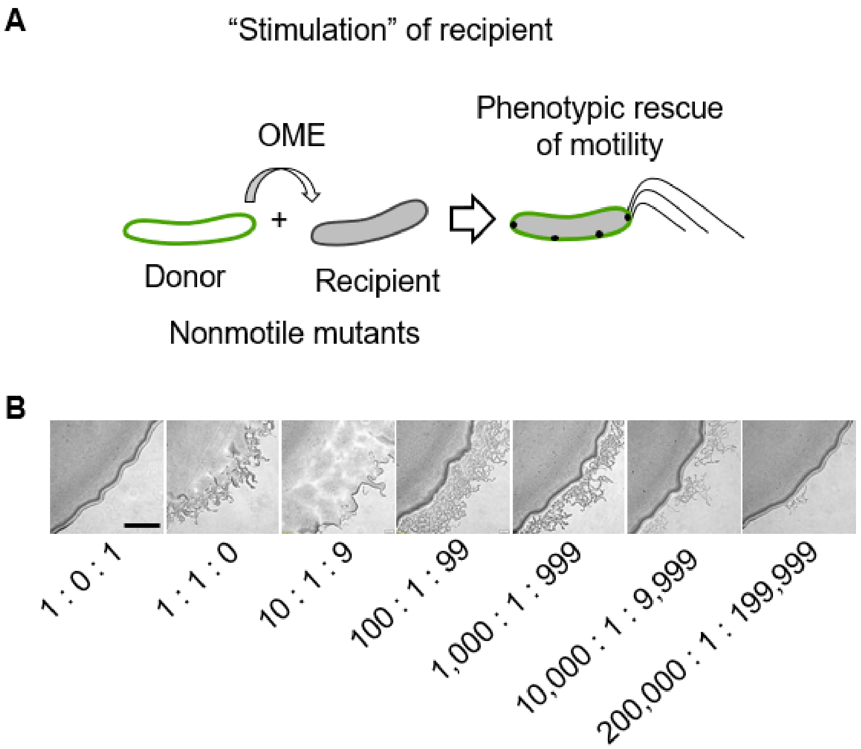


**Fig. S3** Outline and sensitivity of library screen. **A**) Schematic representation of the stimulation assay. Recipient contains two mutations in outer membrane motility lipoproteins (Δ*cglC* Δ*tgl*) required for A- and S-motility, respectively. The nonmotile donor strain contains these WT proteins (green) and transfers them by OME to transiently restore S-motility (type IV pili) and A-motility (black dots, focal adhesions) to the recipient. **B**) Stimulation assay sensitivity. Donor strain ($traA^{DK805}$) was mixed with the stimulatable recipient strain $traA^{DK805}$, and the non-stimulatable strain $traA^{MCy8401}$ at indicated ratios, to maintain the donor ratio of one half of total cells. See Table S1 for strain details. Micrographs at 24 h; scale bar, 200 μm.

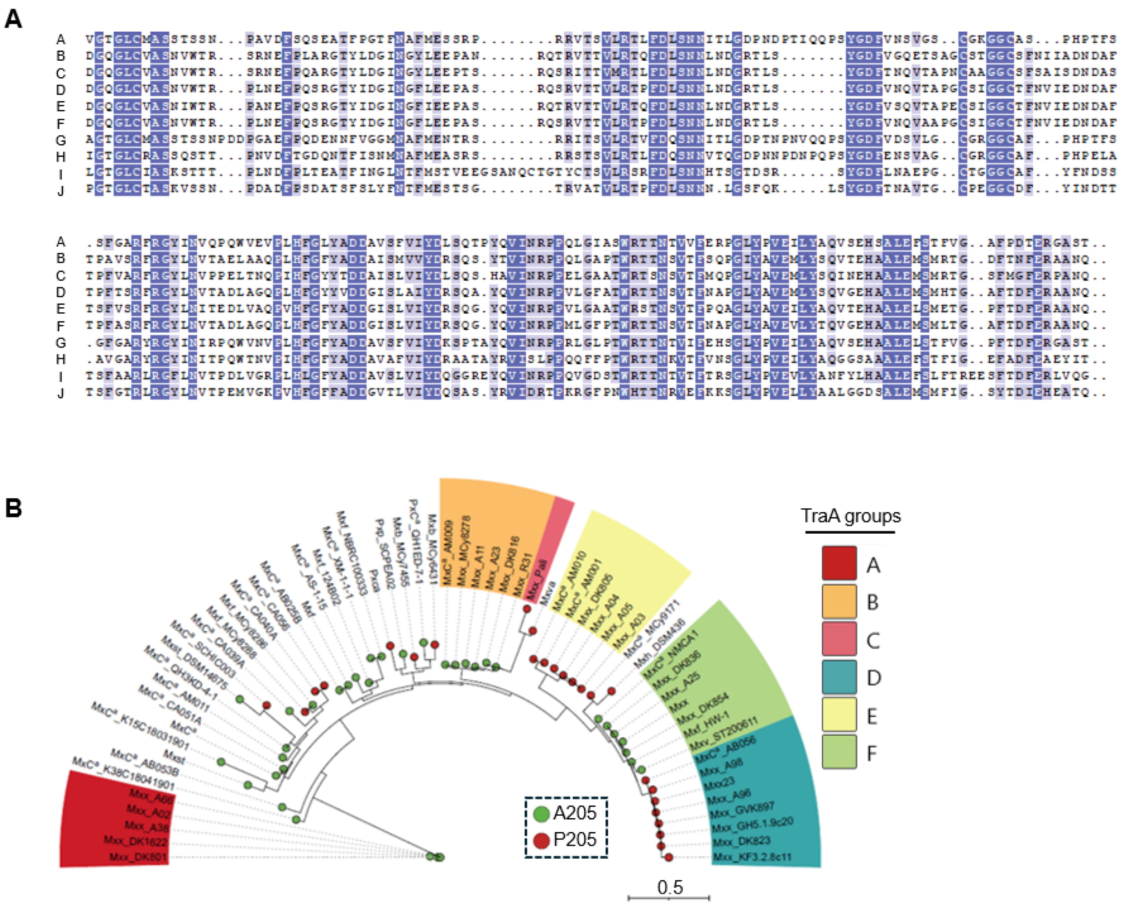


**Fig. S4 A**) Sequence alignment of the VD region from representative TraA group A through J members. Recognition groups labeled on the left. Homologies: Blue, 100%; violet, >75%. **B**) Maximum likelihood tree of the VDs from 62 *Myxococcaceae* TraA orthologs that includes sequences with indels. TraA groups A–F color-coded. Scale bar, number of substitutions per amino acid position.

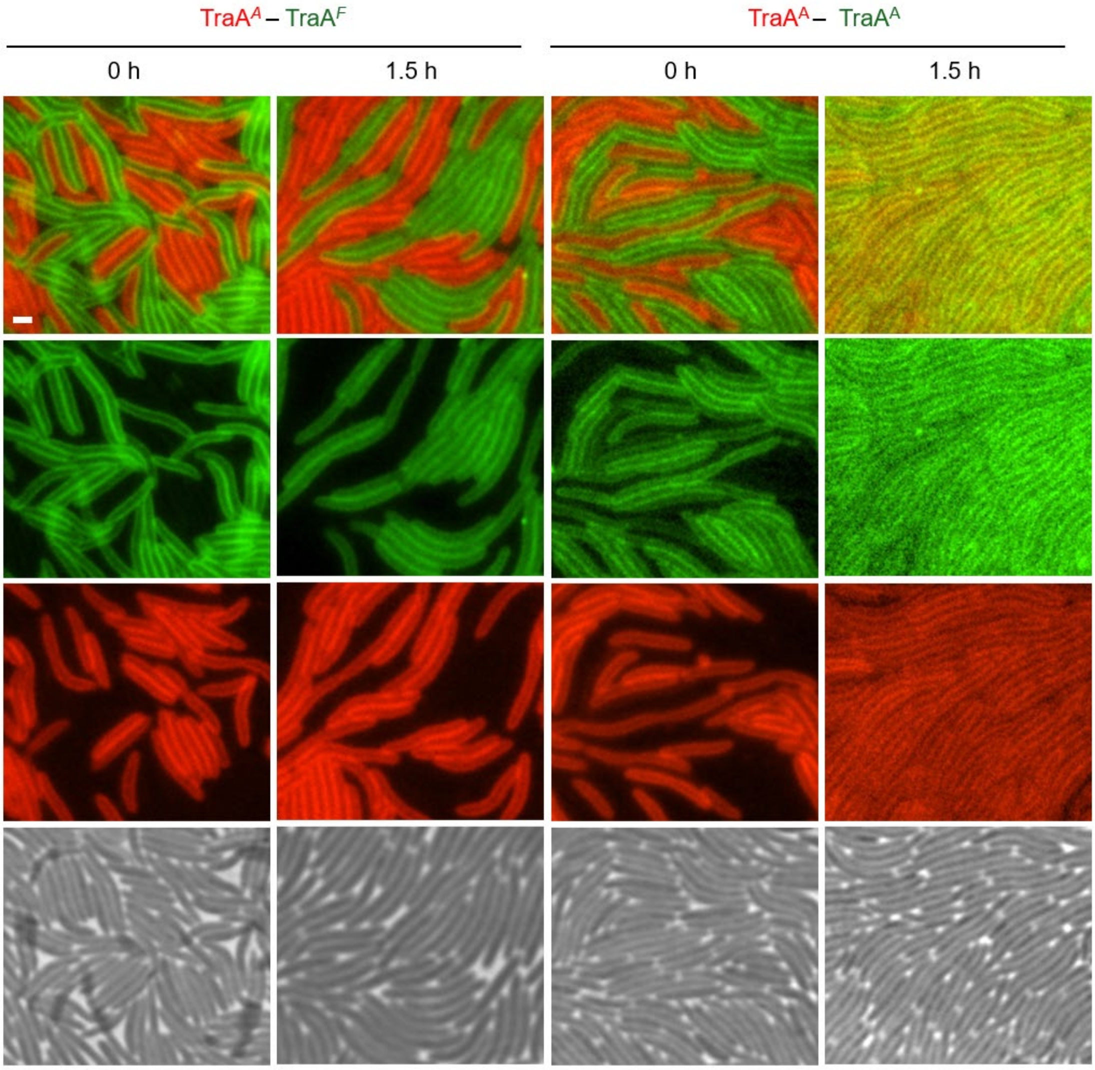


**Fig. S5** Homotypic TraA-dependent OME between cells harboring transferable $SS_{OM}$-GFP or $SS_{OM}$-mCherry reporters. Complementary to Fig. 8A, showing single channel images with merged images. Scale bar, 1 μm.

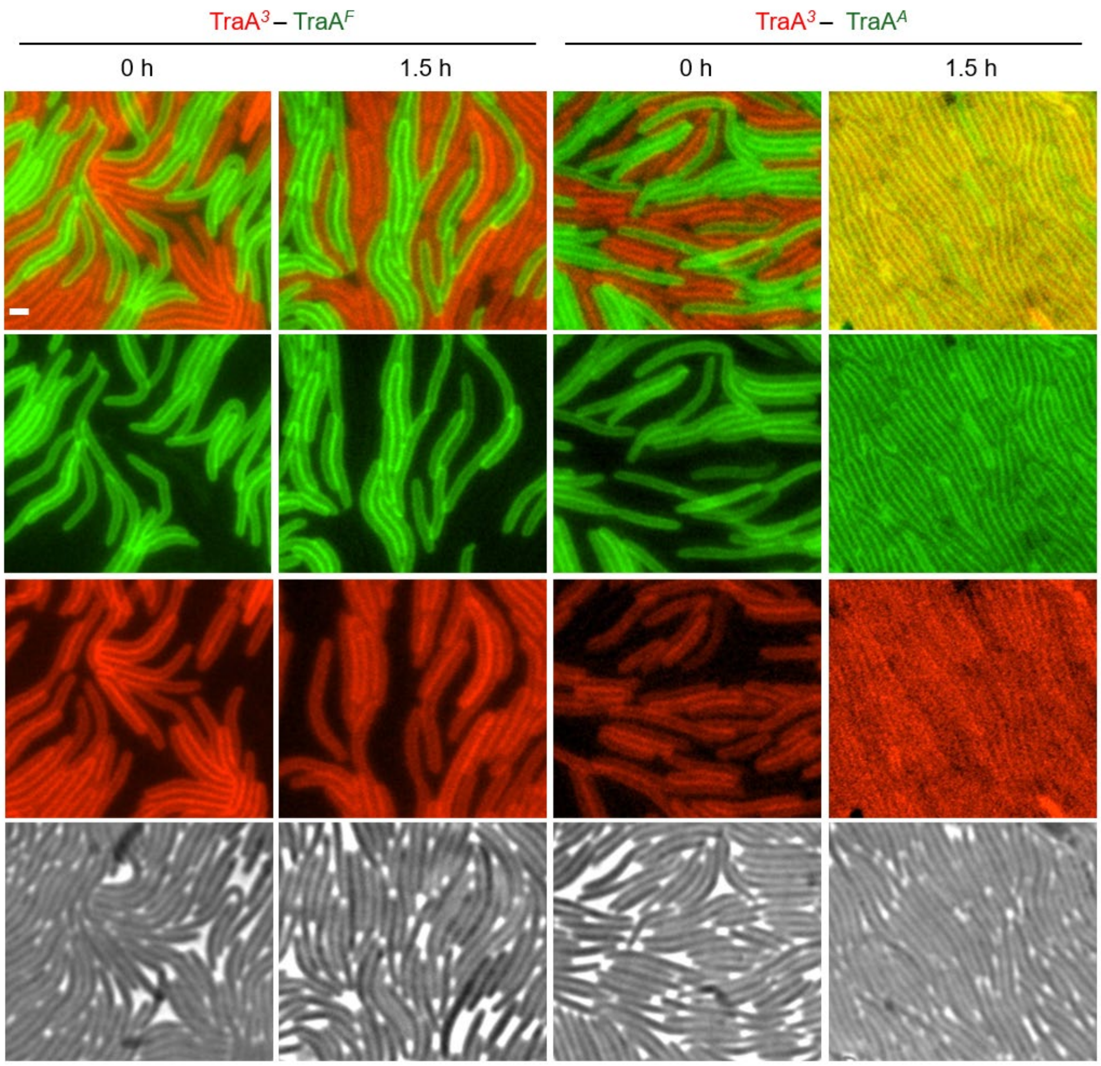


**Fig. S6** Heterotypic TraA-dependent OME between cells harboring transferable $SS_{OM}$-GFP or $SS_{OM}$-mCherry reporters. Complementary to Fig. 8B, showing single channel images with merged images. Scale bar, 1 µm.

**Table S1** Plasmids and strains used in this study

| Plasmids | Relevant features | | Source |
|---|---|---|---|
| pMR3487 | IPTG-inducible promoter, Tc$^R$ | | [1] |
| pΔ*cglC* | Δ*cglC* (deletion cassette) in pBJ114, *galK*, Km$^R$ | | Lotte Søgaard-Andersen |
| pDP28 | Δ*traA* (deletion cassette) in pBJ114, *galK*, Km$^R$ | | [2] |
| pPC16 | P$_{pilA}$-RBSsyn-*traA*$^{DK805}$ in pDP22 (Mx8 *attP*), Km$^R$ | | [3] |
| pPC4 | P$_{pilA}$-RBSsyn-*traAB*$^{Mf}$ in pDP22, Km$^R$ | | [3] |
| pPC5 | P$_{pilA}$-*traA*$^{DK816}$*traB*$^{DK1622}$ in pDP22, Km$^R$ | | [3] |
| pPC26 | P$_{pilA}$-RBSsyn-*traA*$^{MCy5730}$ in pDP22, Km$^R$ | | [4] |
| pDP27 | P$_{pilA}$-RBSsyn-*traA*$^{DK1622}$ in pDP22, Km$^R$ | | [2] |
| pXW6 | P$_{pilA}$-*SS$_{OM}$-mCherry* in pKSAT, Sm$^R$ | | [5] |
| pPC43 | P$_{pilA}$-*SS$_{OM}$-gfp* in pKSAT, Sm$^R$ | | [4] |
| pPC1 | P$_{pilA}$-*SS$_{OM}$*-sfGFP in pKSAT, Sm$^R$ | | [3] |
| pDP21 | P$_{pilA}$-*traA*$^{DK1622}$ in pDP22, Km$^R$ | | [6] |
| pDP22 | pSWU19 Mx8 *attP* cassette, P*pilA*, Km$^R$ | | [2] |
| pDP23 | P$_{pilA}$-*traA*$^{DK816}$ in pDP22, Km$^R$ | | [2] |
| pDP26 | P$_{pilA}$-*traA*$^{Pali}$ in pDP22, Km$^R$ | | [2] |
| pDP24 | P$_{pilA}$-*traA*$^{A96}$ in pDP22, Km$^R$ | | [2] |
| pPC16 | P$_{pilA}$-*traA*$^{DK805}$ in pDP22, Km$^R$ | | [3] |
| pDP25 | P$_{pilA}$-*traA*$^{Mf}$ in pDP22, Km$^R$ | | [2] |
| pPC27 | P$_{pilA}$-*traA*$^{MCy8401}$ in pDP22, Km$^R$ | | [4] |
| pPC28 | P$_{pilA}$-*traA*$^{And48}$ in pDP22, Km$^R$ | | [4] |
| pPC36 | P$_{pilA}$-*traA*$^{MCy8337/DK1622}$ in pDP22, Km$^R$ | | [4] |
| pXW7 | P$_{pilA}$-*traB*,pCR-XL-TOPO; Δ*neoR/kanR*,Mx9 *attP* Zeo$^R$ | | [4] |
| pPC61 | P$_{pilA}$-*traA*$^{Mf(P205A)}$-pSWU19 | | Lab collection |
| pPC62 | P$_{pilA}$-*traA*$^{Mf(P205A)}$-*traB*-pSWU19 | | Lab collection |
| pTG2909 | pMR3487-*traAB*$^{Mf}$, Tc$^R$ | | This study |
| pTG2910 | P$_{pilA}$-RBSsyn-*traA*$^{MCy5730\ (DDPGA \rightarrow LN)}$ in pDP22, Km$^R$ | | This study |
| pTG2911 | P$_{pilA}$-RBSsyn-*traA*$^{DK1622\ (add\ DDP\ before\ AV)}$ in pDP22, Km$^R$ | | This study |
| **Strains** | **Relevant features** | **Experimental use** | **Source** |
| Top10 | *E. coli* cloning strain | Cloning | Lab collection |

| | | | |
|---|---|---|---|
| DK1622 | Wild-type *M. xanthus*, motile | | [7] |
| DK8601 | DK1622 *aglB1* (*aglQ1*) Δ*pilA*::Tc, nonmotile, $Tc^R$ | Fig. 3C and 6E | [8] |
| DW1467 | DK8601 Δ*traA* (markerless), $Tc^R$ | | [2] |
| DK6204 | DK1622 Δ*mglBA* (markerless), nonmotile | | [9] |
| DK8615 | DK1622 Δ*pilQ* (markerless) | | [10] |
| DW1466 | DK1622 *tgl*::Tc Δ*cglC* (markerless), nonmotile, $Tc^R$ | Fig. 2B and 6E | [6] |
| DW2220 | DW1466 Δ*traA* (markerless), $Tc^R$ | | [3] |
| DK101 | *pilQ1* also known as FB or DZF1 vvv | | [10, 11] |
| DK1253 | DK101, *tgl1,* (markerless) | | Lab collection |
| DW2929 | DK1253 Δ*cglC* Δ*traA*, (markerless) | Library strain | This study |
| DW2302 | DK6204 Δ*traA* | Library strain | Lab collection |
| DW1480 | DK1622 Δ*traA* | | [12] |
| DW2930 | DW1480 (pXW6), $Sm^R$ | | This study |
| DW2221 | DW2220 (pDP23), $Km^R$, $Tc^R$ | Fig. 2B | [3] |
| DW2224 | DW2220 (pDP26), $Km^R$, $Tc^R$ | Fig. 2B | [3] |
| DW2222 | DW2220 (pDP24), $Km^R$, $Tc^R$ | Fig. 2B | [3] |
| DW2234 | DW2220 (pPC16), $Km^R$, $Tc^R$ | Fig. 2B | [3] |
| DW2223 | DW2220 (pDP25), $Km^R$, $Tc^R$ | Fig. 2B, 3C and 6B | [3] |
| DW2303 | DW2220 (pPC61), $Km^R$, $Tc^R$ | Fig. 3C and 6B | Lab collection |
| DW2248 | DW2220 (pPC26), $Km^R$, $Tc^R$ | Fig. 6E | [4] |
| DW2931 | DW2302 (pTG2910), $Km^R$ | Fig. 6E | This study |
| DW2932 | DW2220 (pTG2910), $Km^R$ | Fig. 6E | This study |
| DW2933 | DW2302 (pTG2911), $Km^R$ | Fig. 6E | This study |
| DW2934 | DW2220 (pTG2911), $Km^R$, $Tc^R$ | Fig. 6E | This study |
| DW1468 | DW1467 (pDP23), $Km^R$, $Tc^R$ | Fig. 3C | [2] |
| DW1471 | DW1467 (pDP26), $Km^R$, $Tc^R$ | Fig. 3C | [2] |
| DW1469 | DW1467 (pDP24), $Km^R$, $Tc^R$ | Fig. 3C | [2] |
| DW2212 | DW1467 (pPC16), $Km^R$, $Tc^R$ | Fig. 3C | [3] |
| DW1470 | DW1467 (pDP25), $Km^R$, $Tc^R$ | Fig. 3C and 6B | [2] |
| DW2243 | DW1467 (pPC26), $Km^R$, $Tc^R$ | Fig. 6BE | [4] |
| DW2244 | DW1467 (pPC27), $Km^R$, $Tc^R$ | Fig. 6B | [4] |

| | | | |
|---|---|---|---|
| DW2245 | DW1467 (pPC28), Km$^R$, Tc$^R$ | Fig. 6B | [4] |
| DW2262 | DW1467 (pPC36, pXW7), Km$^R$, Tc$^R$, Zeo$^R$ | Fig. 6B | [4] |
| DW2304 | DK8601 Δ*traAB* (pPC62), Km$^R$, Tc$^R$ | | Lab collection |
| DW2935 | DK8615 pXW6 (pMR3487-*traA*$^{3}$**B*), Sm$^R$, Tc$^R$ | | This study |
| DW1463 | DK8601 (pXW6, pDP21), Km$^R$, Tc$^R$, Sm$^R$ | Fig. 7A | [6] |
| DW2201 | DK8601 (pPC1, pDP21), Km$^R$, Tc$^R$, Sm$^R$ | Fig. 8AB and 7AB | [3] |
| DW2287 | DK8601 Δ*traAB* (pPC4, pPC43), Km$^R$, Tc$^R$, Sm$^R$ | Fig. 7A and 8A | [13] |
| DW2936 | DW2929 (pMR3487-*traA*$^{3}$*B*), Tc$^R$ | Fig. 7B | This study |
| DW2937 | DW2303 (pPC43), Km$^R$, Tc$^R$, Sm$^R$ | Fig. 7B | This study |
| DW1478 | DK8615 (pXW6), Sm$^R$ | Fig. 8A | [14] |
| DW2938 | DW2930 (pMR3487-*traA*$^{3}$*B*), Sm$^R$, Tc$^R$ | Fig. 8B | This study |
| DW2939 | DW2304 (pPC43), Km$^R$, Tc$^R$, Sm$^R$ | Fig. 8B | This study |
| DW2940 | DW2302 (pPC16), Km$^R$ | Fig. S3B | This study |
| DW2234 | DW2220 (pPC16), Km$^R$, Tc$^R$ | Fig. S3B | [3] |
| DW2249 | DW2220 (pPC27), Km$^R$, Tc$^R$ | Fig. S3B | [4] |

**Table S2** Primers used in this study

| **Primer name** | **Sequence (5'→3')*** |
|---|---|
| pMR3487-*traAB*$^{Mf}$-XbaI-F | GGATAACAATTAAGGAGGCTCTAGAATGGACGATATCCCTCATTC |
| pMR3487-*traAB*$^{Mf}$-KpnI-R | TGATTACGAAGGCGAGCTCGGTACCCTACGGCTTGGGCGCCGAG |
| pSWU19-F | TGGTAACTGTCAGACCAAG |
| pSWU19-MX8-R | TTCGACGATGGCCTCCAC |
| pSWU19-MX8-F | ACATTGACGTGGAGGCCATC |
| 1622 add DDP before AV-R | AAGTCCACGGCCGGGTCATCCGGATTCGATGATGTGC |
| 1622 add DDP before AV-F | GATGACCCGGCCGTGGACTTCTCTCAG |
| pSWU19-R | AACTTGGTCTGACAGTTAC |
| TraA$^{MCy5730}$-(DDPGA→LN)-R | CTGCGGGAACTCGTTGAGCGGATTCGACGAAGTGCTC |
| TraA$^{MCy5730}$-(DDPGA→LN)-F | CTCAACGAGTTCCCGCAGGACGAA |

Restriction sites underlined.

**Table S3** Abbreviations used to indicated taxa

| **Prefix** | **Species** |
|---|---|
| Mxb | *Myxococcaceae bacterium* |
| Mxf | *Myxococcus fulvus* |
| Mxh | *Myxococcus hansupus* |
| Mxst | *Myxococcus stipitatus* |
| Mxv | *Myxococcus virescens* |
| Mxva | *Myxococcus vastator* |
| Mxx | *Myxococcus xanthus* |
| MxC[a] | *Myxococcus clade* |
| MxC | *Myxococcus* unclassified |
| PxC[a] | *Pyxidicoccus clade* |
| PxCa | *Pyxidicoccus caerfyrddinensis* |
| Pxp | *Pyxidicoccus parkwaysis* |
